# Split suppression gene drives reveal a design tradeoff between inheritance and reproductive load in *Aedes aegypti*

**DOI:** 10.64898/2026.09.17.752470

**Authors:** William A. C. Gendron, Andrea L. Smidler, Ming Li, Tyler Wise, Héctor M. Sánchez C., Jared B. Bennett, Robyn Raban, John M. Marshall, Omar S. Akbari

## Abstract

Mosquito-borne pathogens cause approximately 700,000 deaths annually, highlighting the need for effective and sustainable vector control strategies. CRISPR-based gene drives offer a potential approach for suppressing disease-vector populations, but autonomous drives may spread beyond intended populations. Split gene drives, in which the Cas9 and guide RNA components are genetically separated, provide an inherently confinable alternative because biased inheritance depends on the co-occurrence of both components. Here, we develop the first split suppression gene drives in *Ae. aegypti*, targeting the female reproductive genes *doublesex* (*dsx*) and *yellow-g* (*yg*). The *yg*-targeting drive exhibited moderate super-Mendelian inheritance, reaching 73.4% transmission in the highest-performing cross, whereas the *dsx*-targeting drive exhibited little detectable inheritance bias. Conversely, disruption of *dsx* imposed substantially stronger reproductive fitness costs, including complete sterility in homozygous females, whereas *yg* disruption produced a less severe fertility phenotype. Mathematical modeling parameterized with these experimental data predicted that three of four split-drive configurations could achieve population suppression at lower release intensities or shorter release durations than pgSIT across portions of the modeled parameter space. Together, these results reveal that inheritance efficiency and reproductive load can impose competing constraints on split suppression drive performance: stronger suppression-associated fitness effects do not necessarily translate into improved drive performance when accompanied by limited drive propagation. Our findings establish the feasibility of split suppression gene drives in *Ae. aegypti* and demonstrate that effective suppression drive design requires balancing inheritance efficiency with reproductive fitness costs rather than maximizing either independently.

## Introduction

The yellow fever mosquito, *Ae. aegypti*, is a major vector of arboviruses of global public health importance, including dengue virus (DENV)^1^, chikungunya virus ^2,3^, yellow fever virus ^4^, and Zika virus (ZIKV) ^5,6^. In recent decades, outbreaks of these diseases have caused substantial morbidity and mortality worldwide ^7^, and climate change is expected to expand the geographic range of *Ae. aegypti* and increase the population at risk of arboviral transmission ^8^. Traditional vector control strategies, including insecticides and habitat modification, have had limited success in controlling *Ae. aegypti*, in part because of widespread and increasing insecticide resistance ^9^. Alternative approaches, including Release of Insects carrying a Dominant Lethal gene (RIDL) and *Wolbachia*-based incompatible insect techniques (IIT), have demonstrated the ability to suppress *Ae. aegypti* populations ^10–13^. Self-limiting genetic approaches based on the sterile insect technique (SIT) ^14–17^, including precision-guided sterile insect technique (pgSIT) ^18–20^, are also being developed as promising tools for mosquito population suppression. In particular, pgSIT offers a genetically stable, species-specific, and self-limiting approach that can generate sterile males for targeted population suppression while avoiding the long-term persistence of engineered genetic elements in the environment ^18–20^. The demonstrated potential of these approaches provides a strong foundation for developing complementary genetic control strategies that could further enhance the efficiency, scalability, and durability of mosquito population suppression.

CRISPR-based gene drives provide one potential approach by biasing inheritance of genetic elements above Mendelian frequencies ^21,22^. Homing gene drives (HGDs) have been developed in multiple organisms, including mosquitoes ^23–37^, and can be designed either to suppress populations by disrupting genes required for viability or reproduction or to modify populations by spreading disease-refractory traits. However, autonomous gene drives, in which the components required for drive are genetically linked, have the potential to spread beyond their intended release populations. Split gene drives provide an alternative architecture in which Cas9 and the guide RNA-containing drive element are located at separate genomic loci, with only one component capable of driving ^21,28,33,38–42^. Biased inheritance therefore requires the co-occurrence of both components, and their segregation over successive generations limits sustained drive activity. This architecture provides an inherent mechanism for temporal and potentially spatial confinement, offering greater control over drive persistence than autonomous systems ^21,43^.

Split suppression gene drives have been developed in *D. melanogaster*, *D. suzukii*, and *An. stephensi* ^40,42,44–47^, and split drives for population modification have been developed in *Ae. aegypti* with reduced efficacy^28,33,34,36,48,49^. Autonomous homing gene drives have also been evaluated in *Ae. aegypti* ^50,51^, but split drives designed specifically for population suppression have not. A central challenge for such systems is that the traits responsible for population suppression may themselves compromise drive propagation: disruption of genes with strong effects on reproduction can impose fitness costs on drive-carrying individuals before the drive is efficiently transmitted. We therefore hypothesized that reproductive load and drive propagation could represent competing design considerations, such that targets producing strong reproductive effects would not necessarily produce the greatest overall suppression-drive performance.

Here, we develop the first split suppression gene drives in *Ae. aegypti* by targeting two genes with distinct effects on female reproduction, *yellow-g* (*yg*, AAEL010848) and *doublesex* (*dsx*, AAEL009114). Disruption of *yg* impairs egg viability, whereas disruption of the female-specific *dsx* transcript interferes with female development and reproduction ^27,42,52–58^. We find that the two targets produce contrasting outcomes: *yg* supports moderate super-Mendelian inheritance with comparatively modest reproductive fitness costs, whereas *dsx* produces stronger reproductive defects but little detectable inheritance bias. Because the observed drive performance was modest, we parameterized population models with the experimental data rather than conducting cage suppression trials. Under the modeled release scenarios, three of four split-drive configurations were predicted to achieve population suppression at lower release intensities or shorter release durations than pgSIT across portions of the modeled parameter space. Together, these results establish the feasibility of split suppression drives in *Ae. aegypti* and reveal that reproductive load and inheritance efficiency can impose competing constraints on drive performance, suggesting that effective suppression systems will require balancing these properties rather than maximizing either independently.

## Results

### Design and construction of split suppression drives targeting female reproduction

We developed two split suppression gene drives targeting the *Ae. aegypti* reproductive genes *yellow-g* (*yg*, AAEL010848) and *doublesex* (*dsx*, AAEL009114). Based on previous studies in anopheline mosquitoes ^25,27^, we predicted that disruption of the female-specific exon of *dsx* would impair female development and fertility while preserving male reproductive function. Similarly, disruption of *yg* was expected to reduce egg viability. For each target, we generated a homing-capable gRNA-expressing drive (GDe), designated GDe*_yg_* and GDe*_dsx_* . Each drive expressed a single gRNA from a U6 promoter (U6_AAEL017774)^28^ and contained a 3xP3-tdTomato fluorescent reporter (**Figure 1A; Supplementary Figure 1**). GDe*_yg_* and GDe*_dsx_*were integrated into their respective genomic target loci by homology-directed repair (HDR), and integration was confirmed molecularly **(Supplementary Figure 1)**. Cas9 was supplied from a separate, unlinked transgene to maintain the split-drive architecture. We evaluated two newly generated transgenic lines expressing Cas9 under the control of either the nup50 (AAEL005635) promoter ^26,28^ or the vasa (AAEL004978) promoter ^59,60^. The lines were marked with Opie2-CFP (nup50-Cas9) or Opie2-dsRed (vasa-Cas9) fluorescent reporters and maintained in the wild-type Liverpool genetic background (**Figure 1B and 1C**). Together, these lines enabled evaluation of each GDe with two independent sources of germline Cas9 activity.

**Figure 1.**
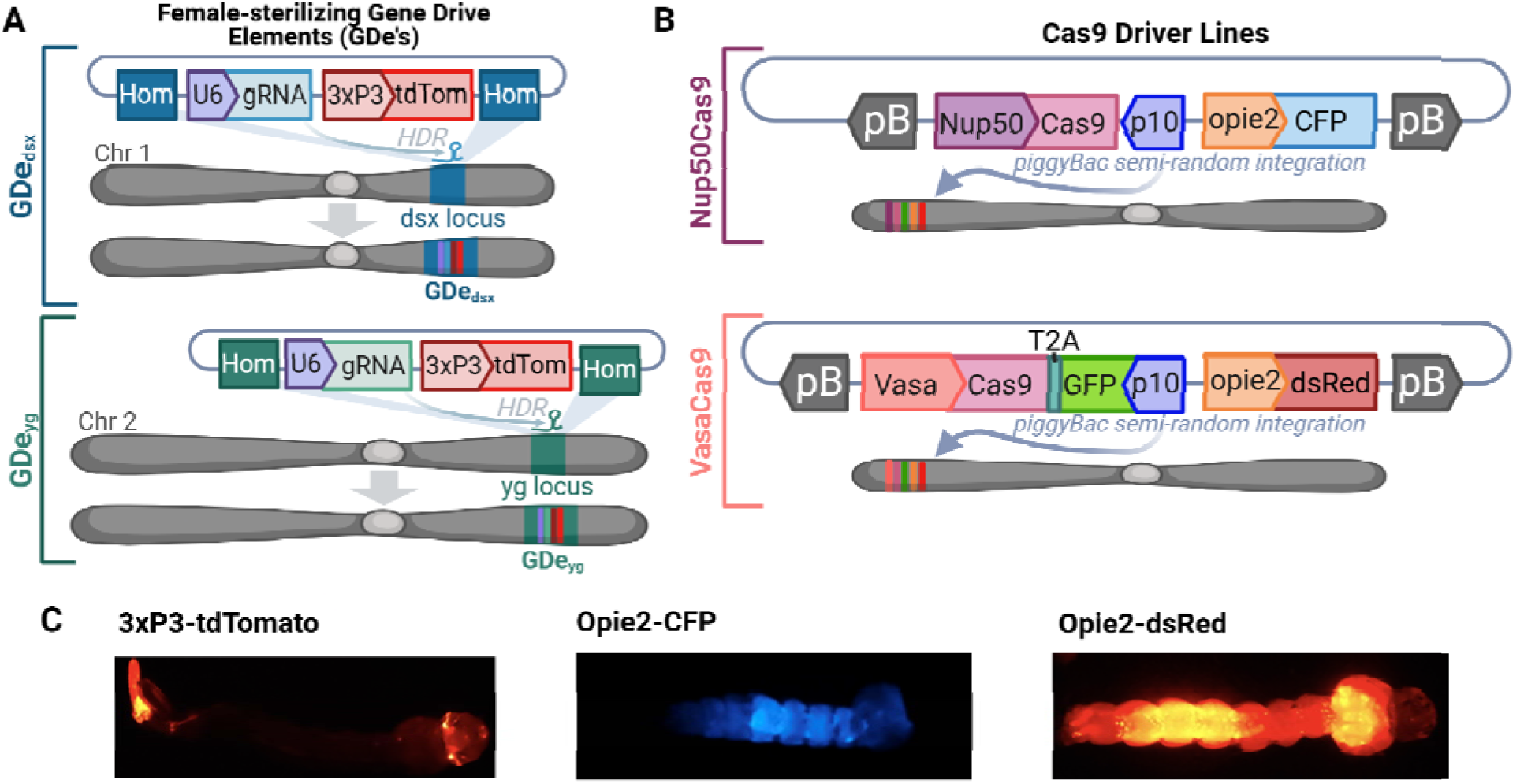
Design of the GDe*_dsx_* and GDe*_yg_* split-drive systems in *Aedes aegypti.* **(A)** Schematic of the GDe*_dsx_*and GDe*_yg_* gene-drive elements (GDe). Each construct contains a U6-driven gRNA targeting either *dsx* or *yg* and a 3xP3-tdTomato fluorescent marker, flanked by homology arms corresponding to the respective genomic target locus. The constructs were integrated by homology-directed repair (HDR) into the *dsx* locus on chromosome 1 or the *yg* locus on chromosome 2. **(B)** Schematic of the Cas9 transgenic constructs used to provide the nuclease component of the split-drive system. Cas9 expression was driven by either the *Nup50* or *vasa* promoter, with Cas9 linked to GFP through a T2A self-cleaving peptide. The Nup50-Cas9 and Vasa-Cas9 constructs were marked with Opie2-CFP and Opie2-dsRed fluorescent reporters, respectively, to facilitate identification of transgenic individuals. **(C)** Representative images of these tracking markers in *Aedes aegypti* larvae.

### GDe*_yg_* and GDe*_dsx_*impose distinct reproductive fitness costs

We first characterized the phenotypic effects associated with each GDe in the absence of Cas9. PCR validated homozygous GDe*_dsx_*/GDe*_dsx_*females were completely infertile (0% egg hatch; n = 21; **Figure 2A**) and exhibited pronounced androgenized morphology, including elongated palps and male-like terminal abdominal structures (**Figure 2B**). In contrast, hemizygous GDe*_dsx_*/+ females appeared morphologically normal but exhibited reduced fertility (66.79% ± 26.43%; n = 77). GDe*_yg_* produced a less severe reproductive phenotype. Homozygous GDe*_yg_*/GDe*_yg_*females retained partial fertility (33.73% ± 26.47%; n = 13), whereas hemizygous GDe*_yg_*/+ individuals exhibited normal fertility (**Figure 2C**). Eggs produced by homozygous GDe*_yg_*females displayed characteristic yellow pigmentation and reduced melanization (**Figure 2D; Supplementary Figure 2**). Adult survival also differed between the two GDe lines. Homozygous GDe*_dsx_* females exhibited significantly reduced survival relative to wild-type females (**Supplementary Figure 3)**, associated with impaired flight and drowning shortly after eclosion. In contrast, homozygous GDe*_yg_* males exhibited a small but statistically significant increase in longevity relative to wild-type males (**Supplementary Figure 3C**), although this effect was not incorporated into subsequent population modeling. Together, these experiments demonstrated substantially stronger reproductive and survival costs associated with GDe*_dsx_* than with GDe*_yg_*.

**Figure 2.**
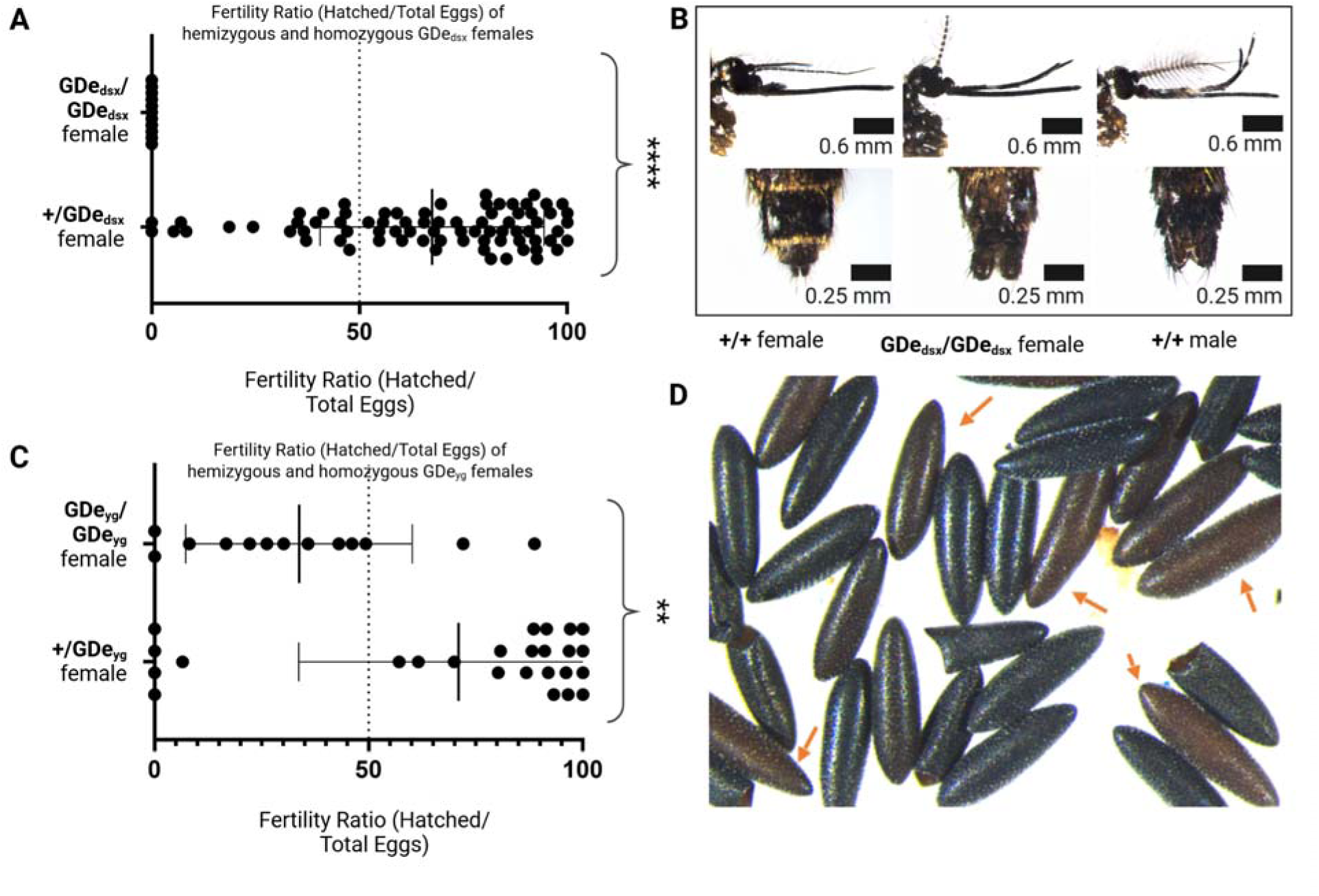
Reproductive and morphological phenotypes associated with GDe*_dsx_*and GDe*_yg_*. **(A)** Fertility of heterozygous (GDe*_dsx_*/+) and homozygous (GDe*_dsx_*/GDe*_dsx_*) females following outcrossing to wild-type (wt) males. Homozygous GDe*_dsx_*females were completely sterile. **(B)** Representative morphology of wt females, homozygous GDedsx females, and wt males. Homozygous GDe*_dsx_* females exhibited androgenized phenotypes, including elongated palps and male-like terminal abdominal structures. **(C)** Fertility of heterozygous (GDe*_yg_*/+) and homozygous (GDe*_yg_*/GDe*_yg_*) females following outcrossing to wt males. Homozygous GDe*_yg_* females exhibited significantly reduced fertility, whereas heterozygous females retained near-wt fertility. **(D)** Representative eggs produced by homozygous GDe*_yg_* females, showing reduced melanization and characteristic yellow pigmentation compared with wt eggs. Fertility was quantified as the proportion of eggs that successfully hatched. Statistical comparisons were performed using Welch’s *t*-test. Data are presented as mean ± SD. *P* < 0.05 (\**), P < 0.01 (**), P < 0.001 (\*\*\**), and *P* < 0.0001 (****).

### GDe*_yg_* but not GDe*_dsx_*exhibits super-Mendelian inheritance

We next evaluated whether the two GDe constructs could achieve biased inheritance when combined with Cas9. In transhemizygous mosquitoes {GDe/+; Cas9/+}, Cas9-mediated cleavage of the wild-type target allele in the germline can be repaired by HDR using the GDe-containing homolog as a template, thereby increasing GDe transmission above the Mendelian expectation of 50%. hemizygous GDe mosquitoes were crossed with either the Nup50Cas9 or VasaCas9 line to generate G1 transhemizygotes, which were subsequently crossed to wild type (**Figure 3A**). Drive activity was quantified by scoring GDe inheritance among the resulting G2 progeny (**Figure 3B; Supplementary Figure 4**). Because parental origin of Cas9 can influence cleavage and homing, crosses involving maternally and paternally inherited Cas9 were analyzed separately and are denoted by “m” and “p,” respectively. As expected, no super-Mendelian inheritance was observed in the absence of Cas9 (**Supplementary Figure 4**). GDe*_dsx_* exhibited little evidence of biased inheritance with either Cas9 line, irrespective of Cas9 parental origin (**Figure 3B**). Transmission was generally at or below the expected Mendelian frequency, with the highest observed GDe*_dsx_* inheritance of 57.64% ± 12.63% in male progeny from crosses involving maternally inherited Nup50Cas9. In contrast, GDe*_yg_* exhibited biased inheritance across Cas9 combinations. The strongest transmission was observed in male progeny from GDe*_yg_* crosses involving maternally inherited VasaCas9, in which the GDe was inherited by 73.40% ± 15.59% of offspring, significantly exceeding Mendelian expectations (**Figure 3B**). TIDE analysis of the *yg* target locus in the Nup50Cas9/GDe*_yg_* cross detected approximately 72.1% sequence disruption (**Supplementary Figure 5A and 5B**), indicating substantial Cas9-mediated cleavage at the target locus. Target-site disruption in the Nup50Cas9/GDe*_dsx_* cross was also clearly detected by sequencing when compared to GDe*_dsx_* crossed to wildtype mosquitoes with 46.6% mutation activity (**Supplementary Figure 5C and 5D**). Thus, despite the stronger reproductive phenotype associated with GDe*_dsx_* , GDe*_yg_* exhibited substantially greater inheritance bias.

**Figure 3.**
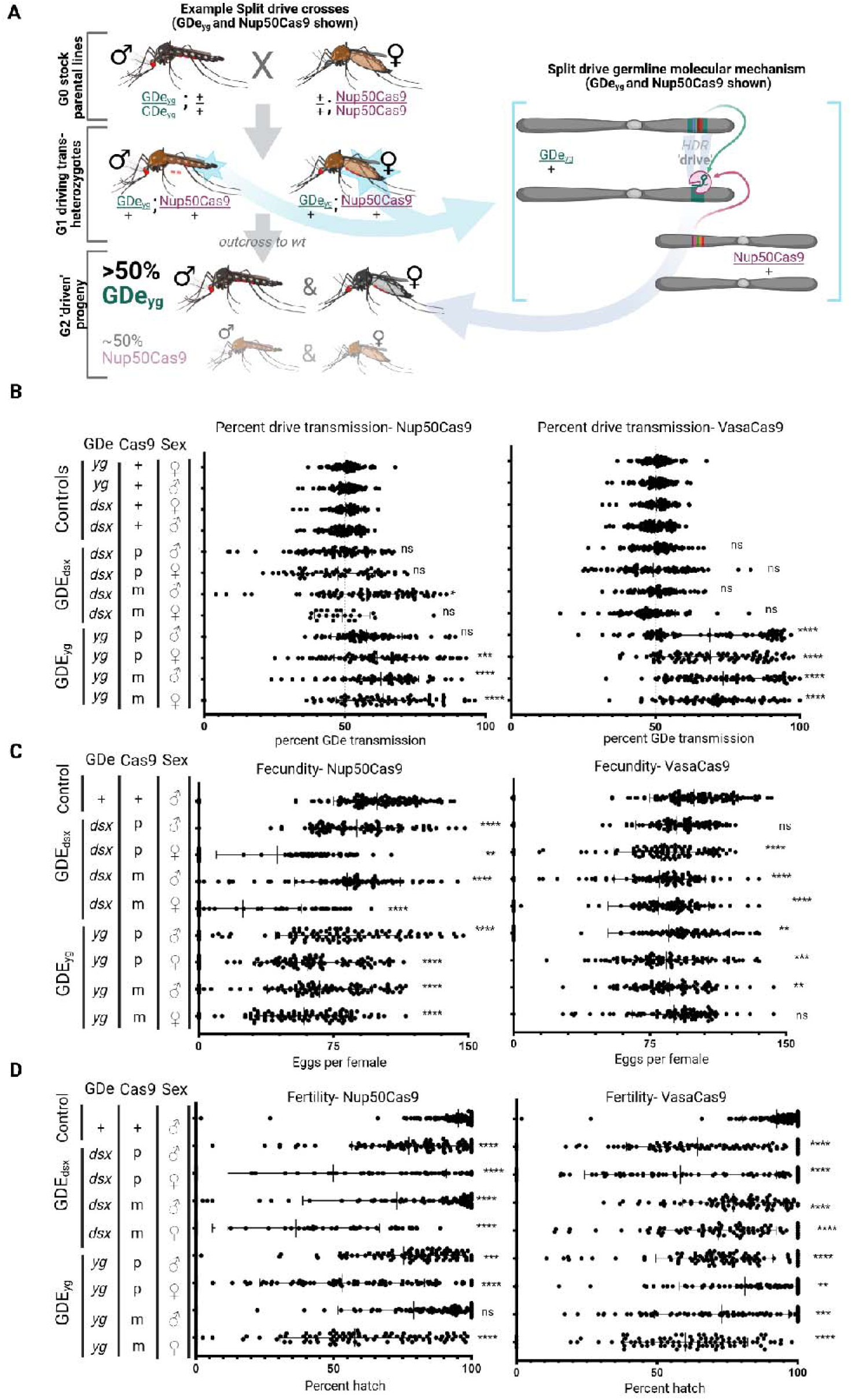
Split-drive inheritance and reproductive fitness effects in *Aedes aegypti*. **(A)** Crossing scheme used to evaluate split-drive activity. Heterozygous GDe*_yg_*/+ or GDe*_dsx_*/+ mosquitoes were crossed with homozygous Nup50-Cas9 or Vasa-Cas9 mosquitoes to generate G1 transheterozygous progeny carrying both the GDe and Cas9 components. G1 transheterozygous males and females were subsequently crossed to wild-type (wt) mosquitoes to assess GDe inheritance and reproductive fitness. **(B)** GDe inheritance frequencies among G2 progeny. Bars indicate the proportion of offspring inheriting GDe*_yg_* or GDe*_dsx_*; the dashed line indicates the expected Mendelian inheritance frequency of 50%. **(C)** Female fecundity, quantified as the number of eggs laid per female following outcrossing to wt males. **(D)** Female fertility, quantified as the proportion of eggs that successfully hatched. Maternal and paternal inheritance of the Cas9 transgene are denoted by “m” and “p,” respectively; “+” denotes the wt allele, and sex symbols indicate the sex of the G1 transheterozygous parent actively driving in the germline. Statistical comparisons of GDe inheritance were performed relative to the corresponding GDe-only controls, whereas fecundity and fertility were compared with the corresponding wt controls using one-way ANOVA followed by Tukey’s multiple-comparisons test. Data are presented as mean ± SD. *P*< 0.05 (\**), P < 0.01 (**), P < 0.001 (\*\*\**), and *P* < 0.0001 (****).

### Split-drive configurations reduce female fecundity and fertility

We next quantified the reproductive effects associated with the active split-drive combinations by measuring female fecundity and fertility in progeny derived from crosses containing both a GDe and Cas9. Fecundity was reduced relative to wild-type controls across the evaluated crosses (**Figure 3C**). The strongest reduction was observed in females from GDe*_dsx_* crosses involving maternally inherited Nup50Cas9, which produced an average of 24.73 ± 32.31 eggs per female. Reduced fecundity was observed with both Nup50Cas9 and VasaCas9, with generally stronger effects in the Nup50Cas9 background. Fertility also varied substantially among split-drive combinations, ranging from approximately 38% to 83% egg hatch (**Figure 3D**). The lowest fertility was observed in females from GDe*_dsx_* crosses involving maternally inherited Nup50Cas9, which exhibited a mean fertility of 38.73% ± 27.86%. GDe*_yg_* crosses also reduced fertility, with the strongest effects observed in combination with Nup50Cas9 regardless of Cas9 parental origin (58.96% ± 26.44% and 58.38% ± 26.28%, respectively). Reproductive costs depended on both the target locus and Cas9 driver, with the strongest effects in the GDe*_dsx_*/Nup50-Cas9 configuration.

### Cas9-associated disruption of *dsx* produces androgenized female phenotypes

To further characterize the reproductive defects associated with GDe*_dsx_*, we examined visible morphological phenotypes in females, including changes in the head, palps, and terminal abdominal structures (**Figure 2B**). Female offspring were classified as exhibiting weak or strong androgenization based on palp morphology (**Supplementary Figure 6**). These phenotypes were detected in homozygous GDe*_dsx_* females and in offspring derived from crosses combining GDe*_dsx_*, with Cas9. The severity of androgenization was associated with impaired reproductive behaviors. Females exhibiting stronger androgenized phenotypes were less likely to fly successfully, blood feed, or oviposit (**Supplementary Figure 7**), all of which are required for successful reproduction. The occurrence of these phenotypes in GDe*_dsx_*/Cas9 crosses is consistent with Cas9-mediated disruption of the *dsx* locus despite the absence of substantial GDe*_dsx_* inheritance bias. The occurrence of these phenotypes, together with sequencing evidence of target-site disruption (**Supplementary Figure 7**), confirms that Cas9 and the GDe*_dsx_* gRNA were functional despite the absence of substantial inheritance bias.

### Experimentally parameterized models predict population suppression by selected split-drive configurations

Because the experimentally measured inheritance bias was modest, we used the MGDrivE simulation framework ^61^ to investigate the potential population-level performance of the split suppression systems rather than conducting cage suppression trials. Simulations considered a randomly-mixing population of 10,000 adult *Ae. aegypti* and compared split-drive releases with pgSIT, a self-limiting genetic suppression system ^18–20,62–64^. To facilitate comparison between technologies, releases were modeled as weekly releases of adult males. Population and construct parameters are provided in **Supplementary Table 1**, and simulations were run for up to 10 years to quantify the duration of population suppression. We subsequently parameterized the *yg* and *dsx* split-drive systems using the experimentally-measured inheritance, fecundity, and fertility values and simulated releases across a range of weekly release ratios and durations (**Figure 4**). Releases were simulated for up to 30 consecutive weeks, with only adult males released. Release ratios ranged from 1-100 transgenic males per wild female. Suppression was evaluated using the “window of protection” metric, defined as the duration for which the non-transgenic female population remained at least 90% below its pre-release level in at least 50% of simulations. Under these modeled conditions, three of the four evaluated split-drive configurations achieved longer windows of protection than pgSIT across portions of the release parameter space: GDe*_yg_*/Nup50Cas9, GDe*_yg_*/VasaCas9, and GDe*_dsx_*/Nup50Cas9 were predicted to suppress populations at lower release intensities or durations than pgSIT, whereas GDe*_dsx_*/VasaCas9 showed little suppression activity (**Figure 4C**).

**Figure 4.**
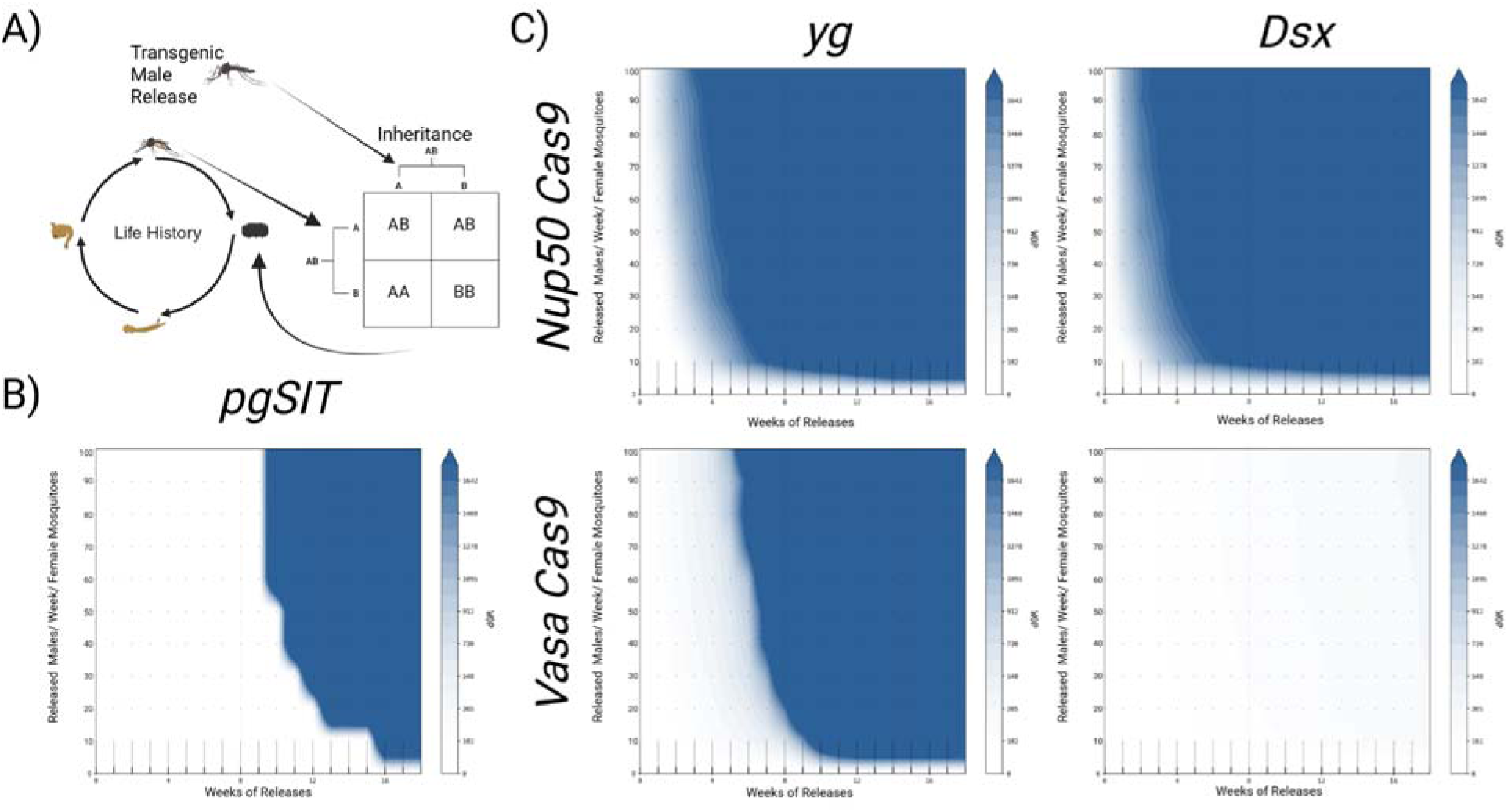
Modeled population suppression by split-drive systems relative to pgSIT. **(A)** Overview of the MGDrivE simulation framework used to evaluate population suppression by the split-drive systems. Simulations incorporated experimentally determined inheritance and fitness parameters for the GDe*_yg_*and GDe*_dsx_* systems, together with shared *Aedes aegypti* life-history parameters. **(B)** Modeled population suppression by precision-guided sterile insect technique (pgSIT), used as a non-driving suppression comparator. Simulations were initialized with a randomly mixing population of 10,000 adult *Ae. aegypti* at demographic equilibrium and modeled weekly releases of transgenic males across a range of release ratios and durations. The x-axis indicates the number of consecutive weekly releases, and the y-axis indicates the number of released transgenic males per wild adult female per week. Suppression is expressed as the window of protection (WOP), defined as the duration for which the non-transgenic female population remained at least 90% below its pre-release equilibrium abundance in at least 50% of stochastic simulations. Darker shading indicates a longer WOP, according to the key. **(C)** Modeled suppression for the four experimentally characterized split-drive configurations: GDe*_yg_*/Nup50-Cas9, GDe*_yg_*/Vasa-Cas9, GDe*_dsx_*/Nup50-Cas9, and GDe*_dsx_*/Vasa-Cas9. Split-drive simulations were parameterized using the experimentally measured inheritance, fecundity, and fertility values reported in this study and were evaluated across the same release ratios and durations used for the pgSIT comparison. Shared bionomic and demographic parameters are provided in **Supplementary Table 1**.

We further determined the minimum release ratios required to achieve suppression following 8, 12, 16, and 20 consecutive weekly releases (**Supplementary Tables 2–5; Supplementary Figure 8**). Under the modeled conditions, pgSIT required release ratios of approximately 150:1, 20:1, and 9.5:1 for 8, 12, and 16-week release programs, respectively (**Supplementary Figure 9**). These simulations predict that selected split-drive configurations could achieve population suppression with lower modeled release ratios or shorter release durations than pgSIT under specific parameter regimes. Finally, we investigated whether combining the inheritance characteristics observed for GDe*_yg_*with the stronger suppression-associated fitness effects of GDe*_dsx_*would enhance population suppression. An idealized system combining these properties did not substantially increase suppression relative to the experimentally characterized drives (**Supplementary Figure 10**). Thus, increasing the magnitude of reproductive fitness effects did not necessarily improve population-level suppression under the modeled conditions. Together, the experimental and modeling results revealed contrasting relationships between reproductive fitness effects and drive performance. GDe*_dsx_*produced severe reproductive phenotypes, including complete infertility in homozygous females, but exhibited little detectable inheritance bias. In contrast, GDe*_yg_* imposed less severe reproductive fitness costs but achieved substantially greater super-Mendelian inheritance. Population modeling further indicated that simply combining stronger reproductive effects with improved inheritance did not necessarily enhance suppression under the evaluated conditions. These findings demonstrate that the magnitude of a suppression phenotype alone is not sufficient to predict the performance of a suppression gene drive. Instead, population-level performance depends on the balance between reproductive load and the ability of the drive to propagate through the population.

## Discussion

Here, we developed the first split suppression gene drives in *Ae. aegypti* and identified an important design tradeoff between reproductive load and drive propagation. GDe*_dsx_* produced strong reproductive effects, including complete infertility in homozygous females, but exhibited little detectable inheritance bias. In contrast, GDe*_yg_* imposed more moderate reproductive costs while achieving super-Mendelian inheritance, reaching 73.4% transmission in the highest-performing cross. These contrasting outcomes suggest that the strongest suppression phenotype does not necessarily produce the most effective suppression drive. Instead, drive performance depends on balancing the reproductive consequences of target disruption with sufficient inheritance to propagate those effects through a population.

Because GDe*_dsx_* and GDe*_yg_* target distinct genomic loci with different gRNAs, their contrasting inheritance efficiencies should not be interpreted as evidence for a general inverse relationship between reproductive load and homing efficiency. Rather, locus-specific differences in cleavage and DNA repair outcomes are also likely to contribute. Importantly, the limited inheritance bias observed for GDe*_dsx_*did not reflect an absence of target cleavage. Cas9-mediated disruption of *dsx* was supported by both sequencing and the highly penetrant androgenized phenotype. Similarly, TIDE analysis of GDe*_yg_*/Nup50Cas9 detected approximately 72.1% target-site disruption despite only moderate inheritance bias. These observations highlight an important distinction between target cleavage and productive homing: efficient cleavage does not necessarily result in HDR-mediated copying of the drive element. Previously, we developed a split drive targeting the white locus using a similar Nup50Cas9 line that achieved inheritance rates of up to 80.5 ± 5.0%, further illustrating that drive performance can vary substantially among genomic targets even when similar Cas9 expression architectures are used ^28^. Direct sequencing of repair products will therefore be important for determining why efficient target disruption at some loci fails to translate into productive homing and for quantifying the formation of potentially cleavage-resistant alleles. The greater drive activity observed with maternally inherited Cas9 further suggests that the timing and parental origin of nuclease activity may influence this balance. Thus, improving drive performance may require not simply increasing Cas9 activity, but optimizing the timing of cleavage to favor HDR-mediated homing over nonproductive repair.

Given the modest inheritance bias observed experimentally, we used population modeling rather than cage suppression trials to evaluate the potential population-level performance of these systems. Despite incomplete drive, three of the four split-drive configurations were predicted to achieve longer windows of protection than pgSIT for specific combinations of release intensity and duration across portions of the modeled parameter space. Notably, GDe*_dsx_*/Nup50Cas9 produced substantial predicted suppression despite little detectable inheritance bias, suggesting that reproductive disruption can contribute meaningfully to population suppression even when homing is inefficient. This result suggests that split suppression systems can derive population-level suppression from both biased inheritance and cleavage-induced reproductive load, with the relative contribution of each depending on drive efficiency and target disruption. In contrast, GDe*_dsx_*/VasaCas9 produced little predicted suppression, indicating that the Cas9 expression background can strongly influence suppression even when inheritance bias is limited. Modeling an idealized system combining the inheritance characteristics of GDe*_yg_* with the stronger reproductive effects associated with GDe*_dsx_* provided limited additional suppression, further illustrating that increasing reproductive load alone does not necessarily improve population-level performance (**Supplementary Figure 10)**. These predictions provide a basis for future evaluation of selected configurations in population cage experiments. Importantly, these comparisons should not be interpreted as indicating that split-drive systems are inherently superior to pgSIT; rather, the two approaches offer distinct operational and biological properties, with pgSIT providing a self-limiting and genetically stable suppression strategy and split drives potentially extending suppression through biased inheritance.

Low homing efficiency remains a major and recurring challenge for gene drives in *Ae. aegypti* ^28,33,34,48–51,65–67^. End-joining repair can compete with the HDR required for homing and generate alleles that either disrupt the target gene or resist further cleavage ^22,33,34,49,68,69^. Future work should therefore focus on defining these repair outcomes while optimizing Cas9 expression, gRNA design, and target selection. Multiplexing may further reduce the formation of resistance alleles ^70^. Because this study evaluated only two suppression targets and two Cas9 expression backgrounds, testing additional targets spanning a broader range of reproductive effects will help define the relationship between reproductive load and inheritance efficiency and determine how broadly the design constraints observed here apply. Environmental context may also influence suppression phenotypes; for example, *yg* has been implicated in egg desiccation resistance ^55,71^, suggesting that its disruption could impose additional fitness effects under field conditions not captured in laboratory assays.

Split gene drives occupy a useful intermediate position between self-limiting suppression technologies and fully autonomous gene drives, combining biased inheritance with inherent limitation of drive activity through segregation of the Cas9 and GDe components. Our results establish the feasibility of this approach in *Ae. aegypti* while also defining an important constraint on its design. More broadly, suppression gene drives should be viewed as optimization rather than maximization problems: strong reproductive load may provide limited population-level benefit when drive propagation is inefficient, whereas efficient inheritance may provide little benefit when reproductive effects are insufficient. Effective suppression systems will therefore require simultaneous optimization of inheritance efficiency, reproductive load, and resistance formation.

## Materials and methods

### Mosquito rearing and maintenance

Mosquitoes used in this study were derived from the *Ae. aegypti* Liverpool (LP) strain, which was used as the wild-type (wt) background and from which the reference genome sequence was generated ^72,73^. Mosquitoes were maintained in an Arthropod Biosafety Level 2 insectary at the University of California, San Diego, at 30.0°C and 75% relative humidity under a 12-h light:12-h dark cycle. Eggs were stored for at least four days to allow completion of embryonic development and were subsequently hatched under vacuum. Larvae were fed ground fish food (TetraFin, Germany). Following pupation and adult eclosion, mosquitoes were provided water and 0.3 M aqueous sucrose solution *ad libitum*. Adults were allowed to mate for five days before females were blood-fed on anesthetized *Mus musculus* for 20 min. All animal procedures were approved by the Institutional Animal Care and Use Committee of the University of California, San Diego (protocol S17187).

### Selection of target genes and gRNA sites

To develop the split-drive systems, we targeted two genes involved in female reproduction: *yellow-g* (*yg*), a recessive, somatically expressed female fertility gene ^25^, and the female-specific transcript of *doublesex* (*dsx*). The *Ae. aegypti yg* ortholog is predicted to contribute to eggshell structural integrity, whereas the female-specific *dsx* isoform (AedsxF) is required for female development ^27^. gRNA target sequences within *yg* and *dsx* were identified using CHOPCHOP ^74^ and are listed in **Supplementary Table 6**. To provide germline Cas9 activity, we used *Ae. aegypti* strains expressing Cas9 under germline-active promoters previously shown to support robust target cleavage and efficient homology-directed repair (HDR) ^26,28^ .

### Construct design and assembly

All constructs were assembled using standard molecular biology methods ^75^. GDe constructs were generated following the approach previously described by Li et al ^28^. Briefly, the *piggyBac* backbone was modified to contain homology arms flanking the *yg* or *dsx* target site. Within the homology arms, each GDe contained a U6 promoter driving expression of a target-specific gRNA comprising a 20-bp targeting sequence followed by an 86-bp gRNA scaffold, synthesized as a gBlock (Integrated DNA Technologies [IDT]; **Supplementary Figure 1**). A 3xP3-tdTomato-SV40 fluorescent marker cassette was also positioned between the homology arms to enable identification of GDe-positive individuals. Primers used for construct assembly are listed in **Supplementary Table 6**. To generate the Cas9-expressing lines, Opie2-CFP-SV40 or Opie2-dsRed-SV40 marker cassettes were cloned into *piggyBac* transgenesis plasmids containing Cas9 under the control of either the *vasa* or *nup50* promoter. In both constructs, Cas9 was linked to GFP through a T2A self-cleaving peptide. The resulting transgenes were integrated semi-randomly into the *Ae. aegypti* genome by *piggyBac*-mediated transgenesis. Plasmids were propagated in JM109 chemically competent cells (Zymo Research, T3005) and purified using Zyppy Plasmid Miniprep (Zymo Research, D4037) or Maxiprep (Zymo Research, D4028) kits. Construct sequences were verified by Sanger sequencing (Source BioScience). All plasmids and corresponding sequence maps are available from Addgene (plasmid nos. [241777 (Vector 1066A-pBac-yellow-g gRNA, Homology Arms, tdTomato), 241778 (Vector 1078BpBacdsx gRNA, Homology Arms, tdTomato), 260305 (Vector 874Z-pBac-vasa-Cas9-2a-eGFP-p10-UTR-Opie2-dsRed) and 260439 (Vector 874P1-pBac-AAEL005635(Nup50)-Cas9-p10-UTR-Opie2-CFP)]).

### Microinjection and selection of transformants

Pre-blastoderm 0–3-h-old embryos were used for microinjection as previously described ^26,76,77^. To generate the GDe lines, wild-type (wt) G0 embryos were injected with a mixture containing 100 ng/μL of either the GDe_yg_ or GDe_dsx_ donor plasmid, 100 ng/μL target-specific gRNA, and 100 ng/μL Cas9 protein. Surviving G0 males and females were reciprocally outcrossed to wt mosquitoes. G1 transformants were identified by fluorescence stereomicroscopy (Leica M165FC) based on expression of the corresponding fluorescent markers.

### Molecular confirmation of site-specific integration

To confirm successful integration of the GDe*_yg_* and GDe*_dsx_* cassettes at their respective genomic target loci, genomic DNA was isolated from individual mosquitoes using the DNeasy Blood & Tissue Kit (QIAGEN) according to the manufacturer’s protocol. Site-specific integration was verified by PCR amplification and Sanger sequencing of the genomic integration junctions using primers spanning the construct and adjacent genomic sequences outside the homology arms (**Supplementary Figure 1; Supplementary Table 6**). GDe*_yg_* and GDe*_dsx_*hemizygous and homozygous individuals were further distinguished by PCR using primers flanking the insertion site (**Supplementary Table 6**). The wild-type and GDe-containing alleles produced products of distinct sizes, enabling genotypes to be assigned based on the presence of the wild-type amplicon, the larger GDe-containing amplicon, or both.

### Assessment of split-drive inheritance, reproductive fitness, and target-site editing

To assess split-drive activity, GDe and Cas9 lines were crossed to generate mosquitoes carrying both components of the split-drive system. Specifically, GDe*_yg_*/+ or GDe*_dsx_*/+ mosquitoes were crossed with homozygous Vasa-Cas9 or Nup50-Cas9 mosquitoes to generate G1 transhemizygous progeny carrying both a GDe and Cas9. G1 progeny were identified based on expression of the corresponding fluorescent markers and were subsequently reciprocally crossed to wild-type (wt) mosquitoes to assess GDe transmission and reproductive fitness. Transhemizygous mosquitoes representing each of the four split-drive configurations—{GDe*_yg_*/+; Nup50-Cas9/+}, {GDe*_yg_*/+; Vasa-Cas9/+}, {GDe*_dsx_*/+; Nup50-Cas9/+}, and {GDe*_dsx_*/+; Vasa-Cas9/+}—were crossed to wt mosquitoes for four days. On the fifth day, females were blood-fed on anesthetized mice. Immediately following blood feeding, 30 females from each cross were individually transferred to 50-mL Falcon tubes lined with moist filter paper and allowed to oviposit overnight. Progeny from each female were screened for the 3xP3-tdTomato GDe marker by fluorescence stereomicroscopy, and GDe transmission was calculated as the proportion of progeny inheriting GDe_yg_ or GDe_dsx_. Under Mendelian inheritance, a hemizygous GDe is expected to be transmitted to 50% of offspring; transmission frequencies significantly exceeding 50% were considered evidence of super-Mendelian inheritance (**Figure 3B**). Females that did not blood-feed or failed to produce progeny were excluded from the transmission analysis. Reciprocal crosses were performed to evaluate the effects of sex and parental origin of Cas9 on GDe transmission. Fecundity and fertility were assessed in parallel, with fecundity defined as the number of eggs laid per female and fertility defined as the proportion of eggs that successfully hatched. These assays were used to characterize inheritance and reproductive fitness effects across the four GDe/Cas9 combinations. Additionally, G1 progeny were selected for DNA isolation and PCR amplification of the cleavage site using our TIDE primers for Sanger sequencing and TIDE analysis to confirm Cas9 editing activity **(Supplementary Figure 5, Supplementary Table 6)**.

### Phenotypic and reproductive fitness assays

Phenotypic characterization was performed using a Leica M165FC stereomicroscope equipped with a Leica DMC2900 camera. Pupae and adult mosquitoes were collected separately in Falcon tubes and anesthetized on ice for 5 min before imaging. To assess the reproductive fitness effects associated with the GDe constructs, hemizygous GDe*_yg_*/+ or GDe*_dsx_*/+ males and females were intercrossed to generate hemizygous and homozygous progeny. Genotypes were initially distinguished based on the intensity of the 3xP3-tdTomato fluorescent marker. Nup50-Cas9/+, GDe*_yg_*/+, GDe*_dsx_*/+, and non-fluorescent progeny were maintained as controls, and experimental and control assays were performed concurrently. For fecundity and fertility assays, groups of 35 homozygous or hemizygous males or females were outcrossed to an equal number of wild-type (wt) mosquitoes of the opposite sex and allowed to mate for four days. On the fifth day, females were blood-fed on anesthetized mice. At 48 h post-blood meal, 30 females from each cross were individually transferred to 50-mL tubes lined with moist filter paper and allowed to oviposit overnight. Fecundity was quantified as the number of eggs laid per female, and fertility was calculated as the proportion of eggs that successfully hatched. At least 15 progeny from each cross were screened for fluorescent marker expression using a Leica M165FC stereomicroscope to confirm the expected parental genotypes. For females that failed to produce progeny, genotype was confirmed by PCR using primers 1078B ml1 and 1078B ml4 (**Supplementary Table 2)** to distinguish hemizygous and homozygous individuals. Statistical analyses were performed using GraphPad Prism (GraphPad Software, La Jolla, CA, USA). Differences in GDe transmission, fecundity, and fertility among experimental groups were evaluated using one-way analysis of variance (ANOVA) followed by Tukey’s multiple-comparisons test. For the analyses presented in **Figure 3**, fecundity and fertility were compared with the corresponding wt or parental-line controls, as appropriate. Statistical significance was defined as *P* < 0.05. GDe transmission assays were performed using a similar crossing scheme; however, all larval progeny from each informative cross were screened for the 3xP3-tdTomato marker to determine GDe*_yg_* or GDe*_dsx_* transmission frequencies.

### Adult survival assays

To assess adult longevity, equal numbers of male and female GDe*_yg_*, GDe*_dsx_*, and wild-type (wt) mosquitoes were maintained in small cages and monitored daily for survival (**Supplementary Figure 3**). Statistical analyses were performed using GraphPad Prism 7 (GraphPad Software, La Jolla, CA, USA). Comparisons between hemizygous and homozygous GDe groups were performed using unpaired *t*-tests with Welch’s correction, where appropriate. Survival distributions were compared using the Mantel–Cox (log-rank) test. Differences were considered statistically significant at *P* < 0.05.

### Mathematical modeling

To evaluate the expected population-level performance of the split-suppression drive systems relative to pgSIT, we used the MGDrivE simulation framework ^61^. MGDrivE models egg, larval, pupal, and adult mosquito life stages with overlapping generations, density-dependent larval mortality, and a mating structure in which females retain the genetic material of the male with whom they mate for the duration of their adult lifespan. Full mathematical details of the framework are provided in the S1 Text of Sánchez C. et al. ^61^. For the split-suppression drive simulations, we incorporated effects on fertility, fecundity, and pupation associated with Cas9 inheritance, gRNA inheritance, disruption of the target gene (*dsx* or *yg*), and homing activity, where applicable. Split-drive inheritance and fitness parameters were derived from the experimental measurements reported in this study, whereas fitness parameters for pgSIT were obtained from a previously published study in *Ae. aegypti* ^62^. Bionomic and demographic parameters shared across the modeled systems are provided in **Supplementary Table 1**. For each release scenario, 100 stochastic simulations were performed using a randomly mixing population of 10,000 adult *Ae. aegypti* initialized at demographic equilibrium. Density-independent mortality rates for juvenile life stages were parameterized to be consistent with the population growth rate in the absence of density-dependent mortality, while density-dependent mortality was applied during the larval stage following Deredec et al. ^78^. The suppression performance of the GDe*_yg_* and GDe*_dsx_*split-drive systems was then compared with that of pgSIT under the modeled release scenarios. Population suppression was quantified using the window of protection (WOP) metric, defined as the duration for which the non-transgenic female *Ae. aegypti* population remained at least 90% below its pre-release equilibrium abundance in at least 50% of stochastic simulations. WOP values were evaluated across release intensities and durations, with outcomes following 8, 12, 16, and 20 weeks of releases (for release ratios of 1-100 transgenic males per wild female) summarized in **Supplementary Tables 2–5**.

## Ethics statement

All animal procedures were approved by the Institutional Animal Care and Use Committee of the University of California, San Diego (protocol S17187).

## Acknowledgements

This work was supported by an NIH award (R01AI151004) awarded to O.S.A., and by an NIH award (R01AI190001) awarded to O.S.A. and J.M.M. Modeling support was provided by an NIH award (R01AI189916) awarded to J.M.M. The funders had no role in study design, data collection, analysis, the decision to publish, nor the preparation of the manuscript. We thank Judy Ishikawa for helping to rear all mosquito strains produced in this study.

## Author contributions

O.S.A. conceptualized the study. W.A.C.G. and M.L. performed experiments. H.M.S.C., J.B.B., and J.M.M. performed the mathematical modeling. All authors wrote, edited, and approved the final manuscript.

## Data Availability Statement

All data are provided within this manuscript or the supplemental material. Plasmids used in this study are available for purchase at www.addgene.com [241777 (Vector 1066A-pBac-yellow-g gRNA, Homology Arms, tdTomato), 241778 (Vector 1078B-pBacdsx gRNA, Homology Arms, tdTomato), 260305 (Vector 874Z-pBac-vasa-Cas9-2a-eGFP-p10-UTR-Opie2-dsRed) and 260439 (Vector 874P1-pBac-AAEL005635(Nup50)-Cas9-p10-UTR-Opie2-CFP)].

## Competing Interests

O.S.A. is a founder of Agragene, Inc. Synvect, Inc., and Cloak Biotech with equity interest. The terms of this arrangement have been reviewed and approved by the University of California, San Diego, in accordance with its conflict of interest policies. A.L.S. has a patent on CRISPR-based gene drives (WO2015105928A1). The rest of the authors declare no competing interests.

## Supplementary Figures

**Supplementary Figure 1.**
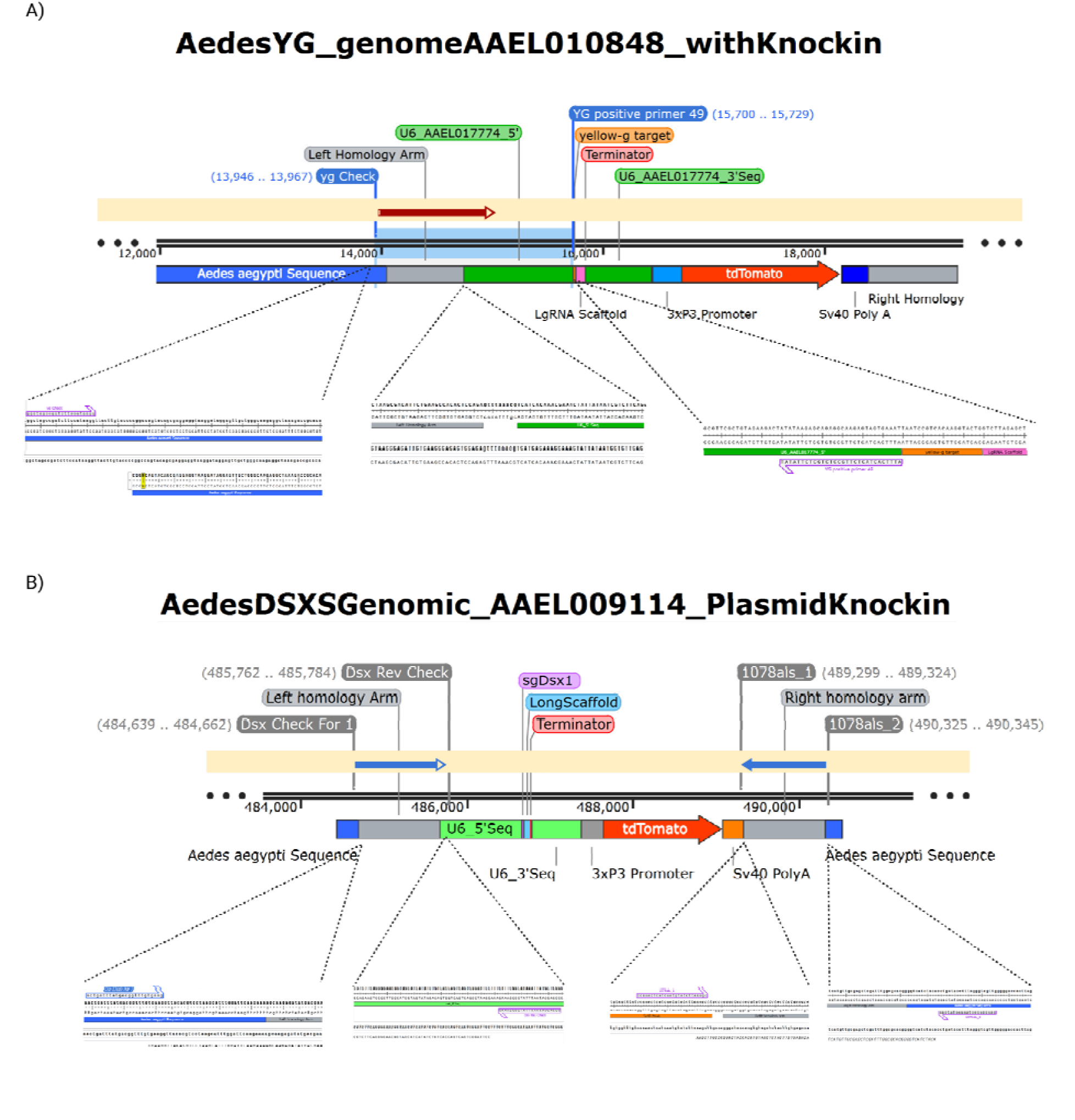
Molecular confirmation of GDe_yg_ and GDe_dsx_ integration. Schemati representations of the gene-drive elements integrated at the Aedes aegypti yellow-g (**A;** GDe_yg_, vector 1066A) and doublesex (**B;** GDe_dsx_, vector 1078B) target loci. The target-specific U6–gRNA cassette, 3×P3–tdTomato fluorescent marker, SV40 polyadenylation signal, and left and right homology arms are indicated. Diagnostic PCR primers positioned within the gene-drive element and in the flanking genomi regions were used to amplify the construct–genome junctions. For GDe_yg_, the left junction was amplified using a primer outside the left homology arm and an internal primer adjacent to the gRNA cassette. For **GDe_dsx_**, both the left and right junctions were amplified using A. aegypti genomic primers outside the corresponding homology arms and construct-specific internal primers. The sequence alignments shown below each schematic confirm the expected construct–genome junctions and site-specific integration at the respective target loci.

**Supplementary Figure 2.**
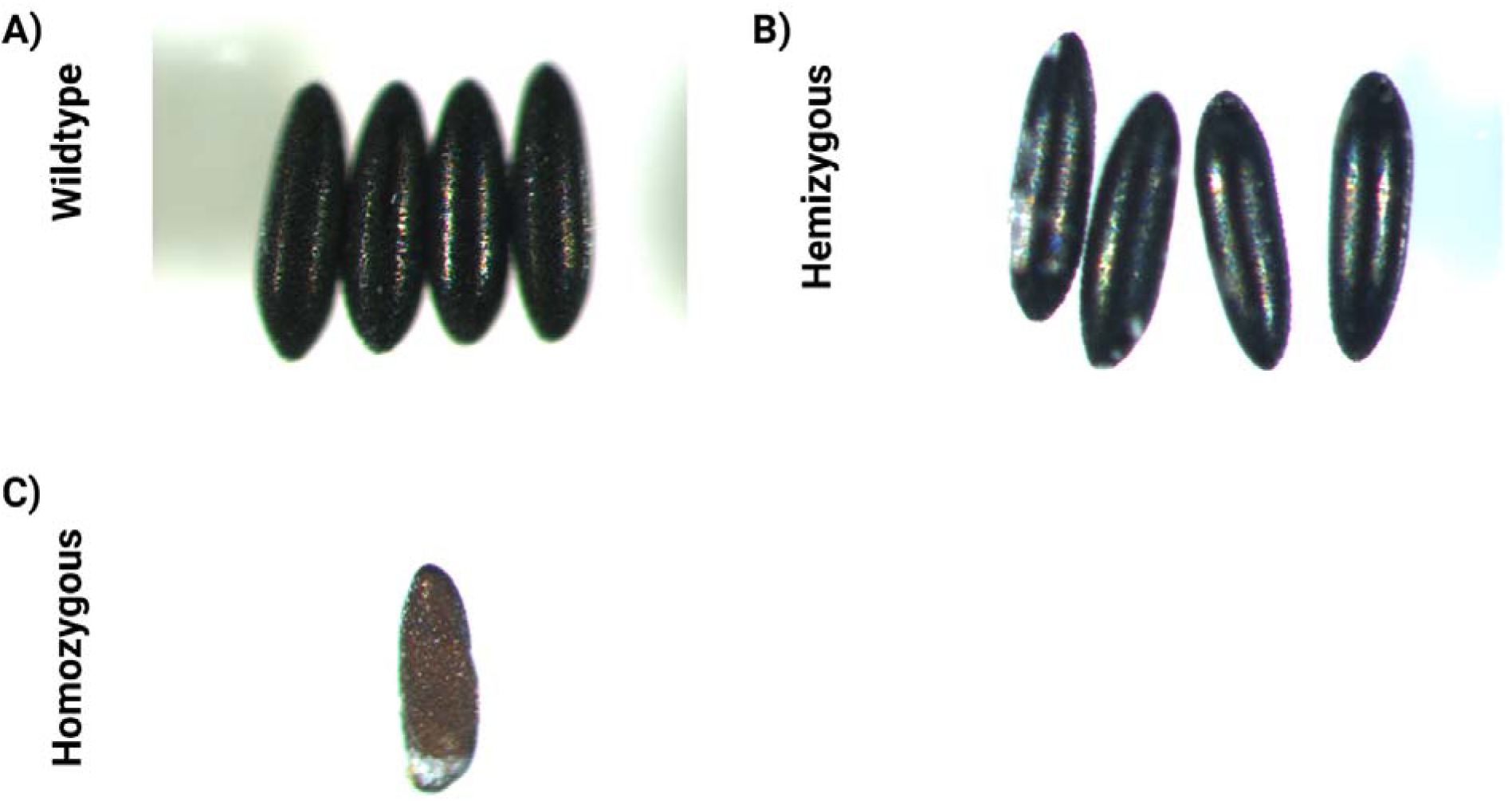
Egg phenotype associated with GDe_yg_. Representative images of eggs produced by **(A)** wild-type (wt), **(B)** heterozygous GDeyg/+, and **(C)** homozygous GDe_yg_/GDe_yg_ females. Homozygous GDe_yg_ females produced eggs displaying altered pigmentation and reduced melanization relative to wild-type and heterozygous controls.

**Supplementary Figure 3.**
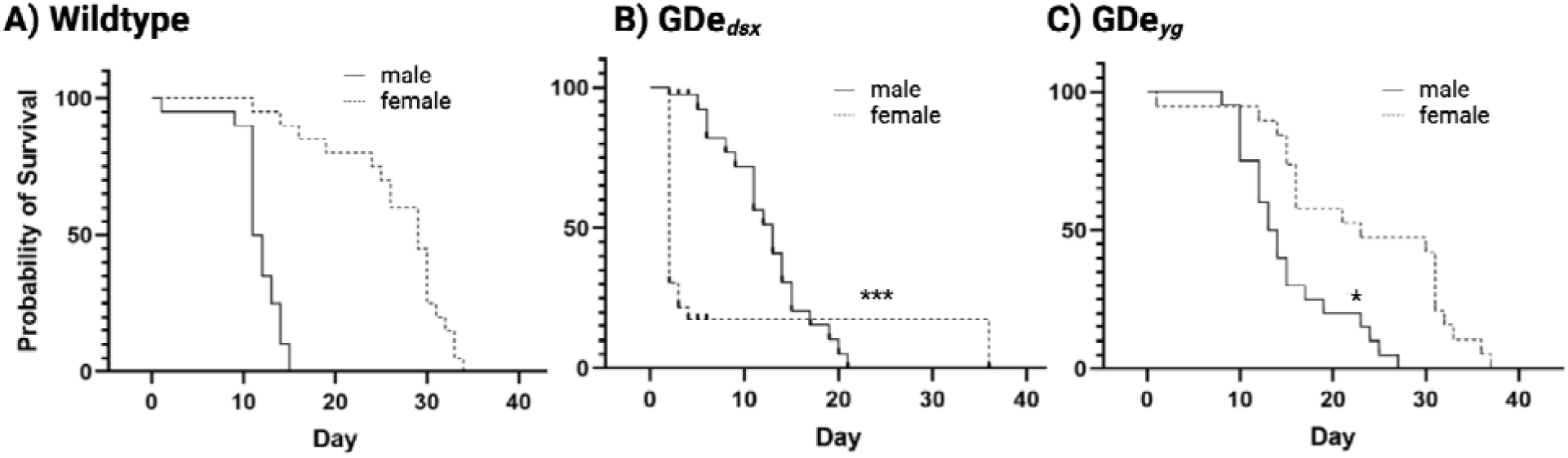
Adult longevity of homozygous GDe_dsx_ and GDe_yg_ mosquitoes. Adult survival was monitored daily in wild-type and homozygous GDe mosquitoes. Survival curves are shown for **(A)**wild-type, **(B)** homozygous GDe_dsx_/GDe_dsx_, and **(C)** homozygous GDe_yg_/GDe_yg_ mosquitoes. For statistical comparisons, homozygous males were compared with wild-type males and homozygous females with wild-type females. Survival distributions were evaluated using the Mantel–Cox log-rank test.

**Supplementary Figure 4.**
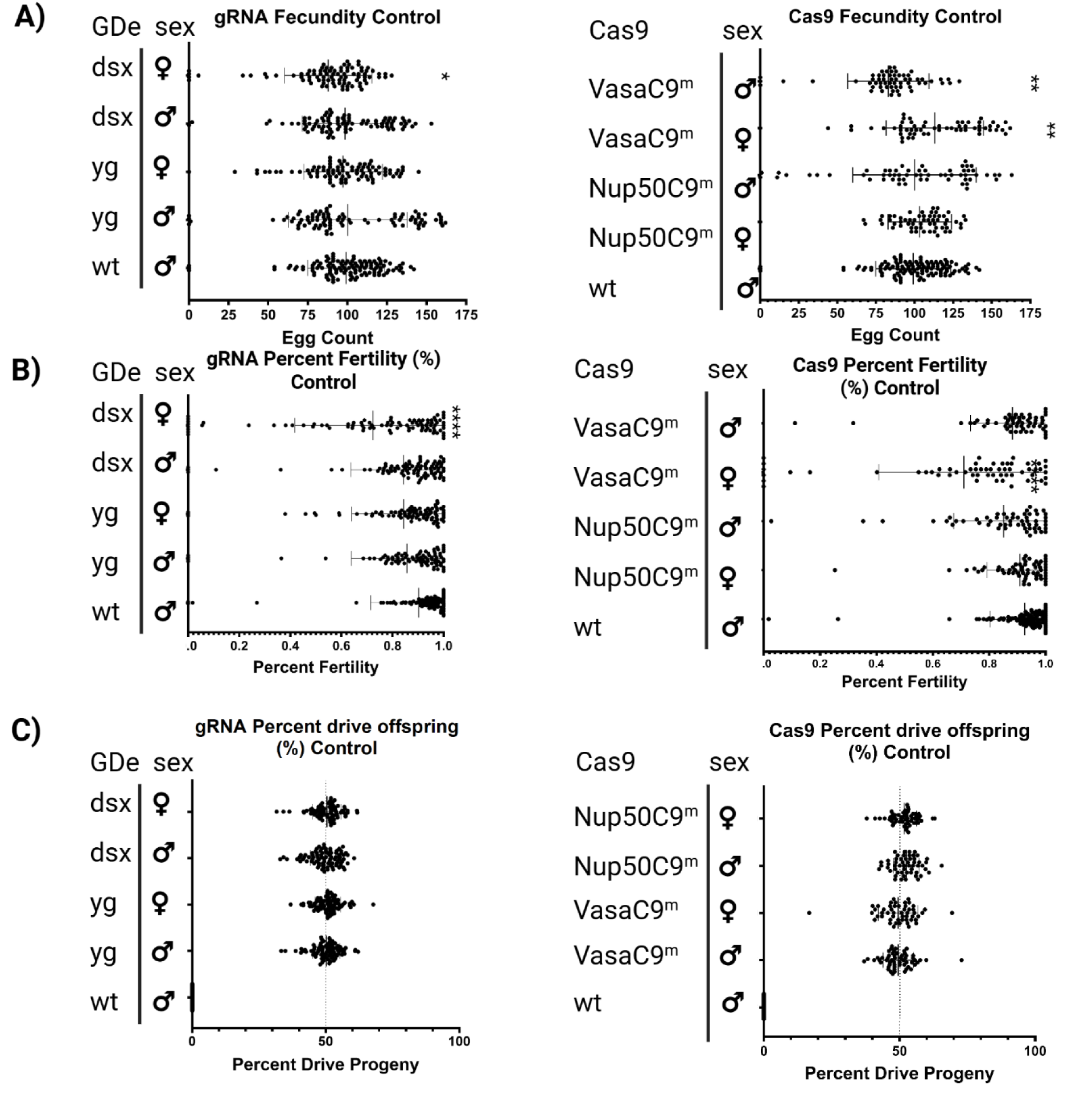
Reproductive fitness and inheritance of the individual GDe and Cas9 parental lines. The individual heterozygous GDe_yg_ and GDe_dsx_ lines and the Cas9-expressing lines were crossed to wild-type mosquitoes to assess baseline **(A)** fecundity, **(B)** fertility, and **(C)** inheritance in the absence of an active split-drive combination. Wild-type controls represent offspring generated from wild-type male × wild-type female crosses. For the Cas9 lines shown, Cas9 was maternally inherited and is therefore denoted with the superscript “m.”

**Supplementary Figure 5.**
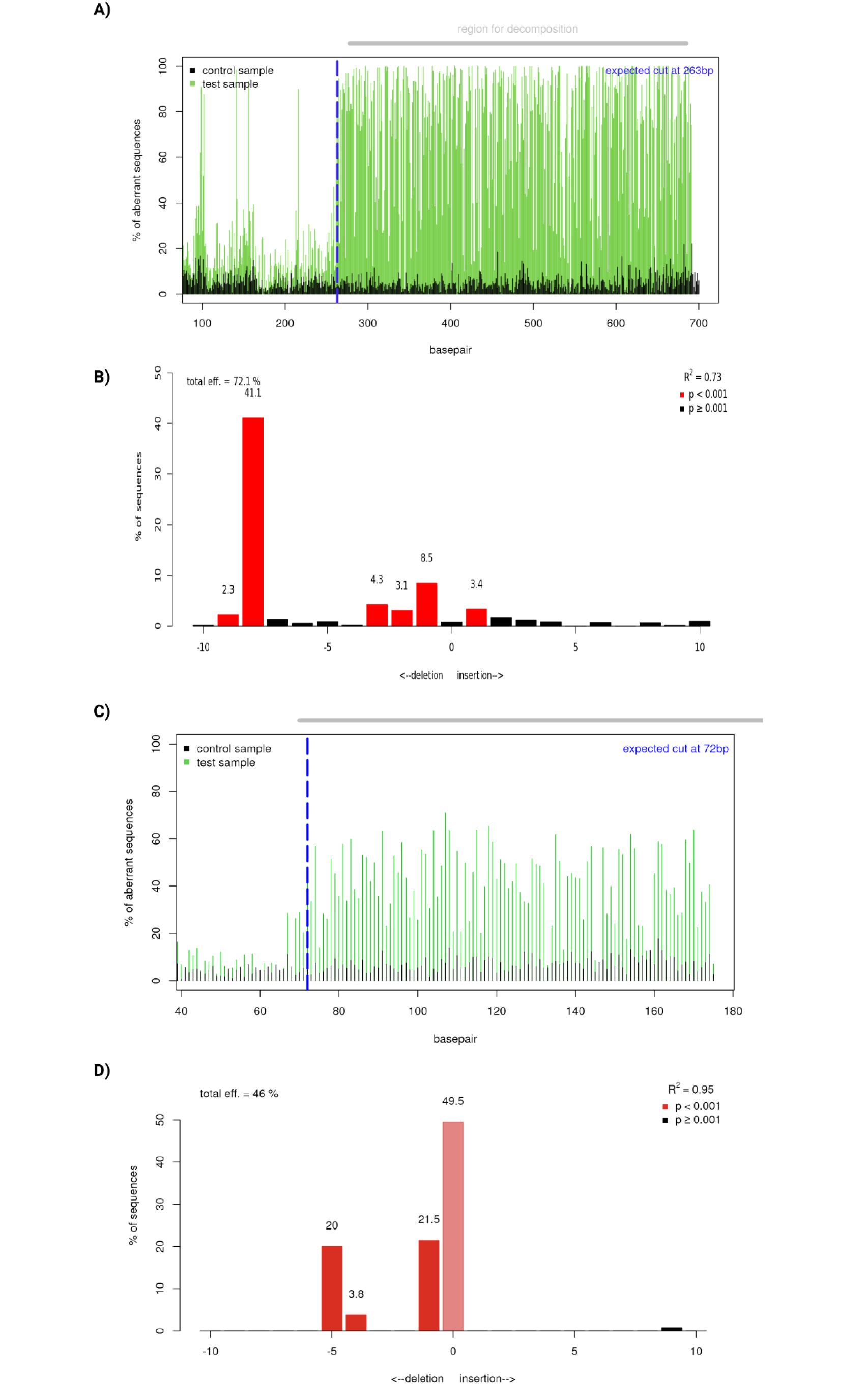
TIDE analysis of Cas9-mediated sequence disruption at the yellow-g and doublesex target sites. Target loci were amplified and analyzed by Sanger sequencing followed by Tracking of Indels by DEcomposition (TIDE). **(A)** Sequence-decomposition plot comparing the yellow-g control sample (GDe_yg_ crossed with wild type; black) with the test sample (GDe_yg_) male × Nup50Cas9 female; green). The blue dashed line marks the expected Cas9 cleavage site, and the gray bar indicates the region used for decomposition. **(B)** Estimated distribution of insertion and deletion (indel) sizes at the yellow-g target site. Negative values indicate deletions, positive values indicate insertions, and zero represents sequences with no change in length. Red bars indicate statistically significant indel classes (p < 0.001), whereas black bars indicate nonsignificant classes (p ≥ 0.001). The estimated overall editing efficiency was 72.1%, with a model fit of (R^2 = 0.73). **(C)** Sequence-decomposition plot comparing the doublesex control sample (GDe_dsx_) crossed with wild type; black) with progeny carrying both GDe_dsx_) and Nup50Cas9 (green). Markings are as described in panel A. **(D)** Estimated indel-size distribution at the doublesex target site, displayed as described in panel B. The estimated overall editing efficiency was 46.0%, with a model fit of (R^2 = 0.95).

**Supplementary Figure 6.**
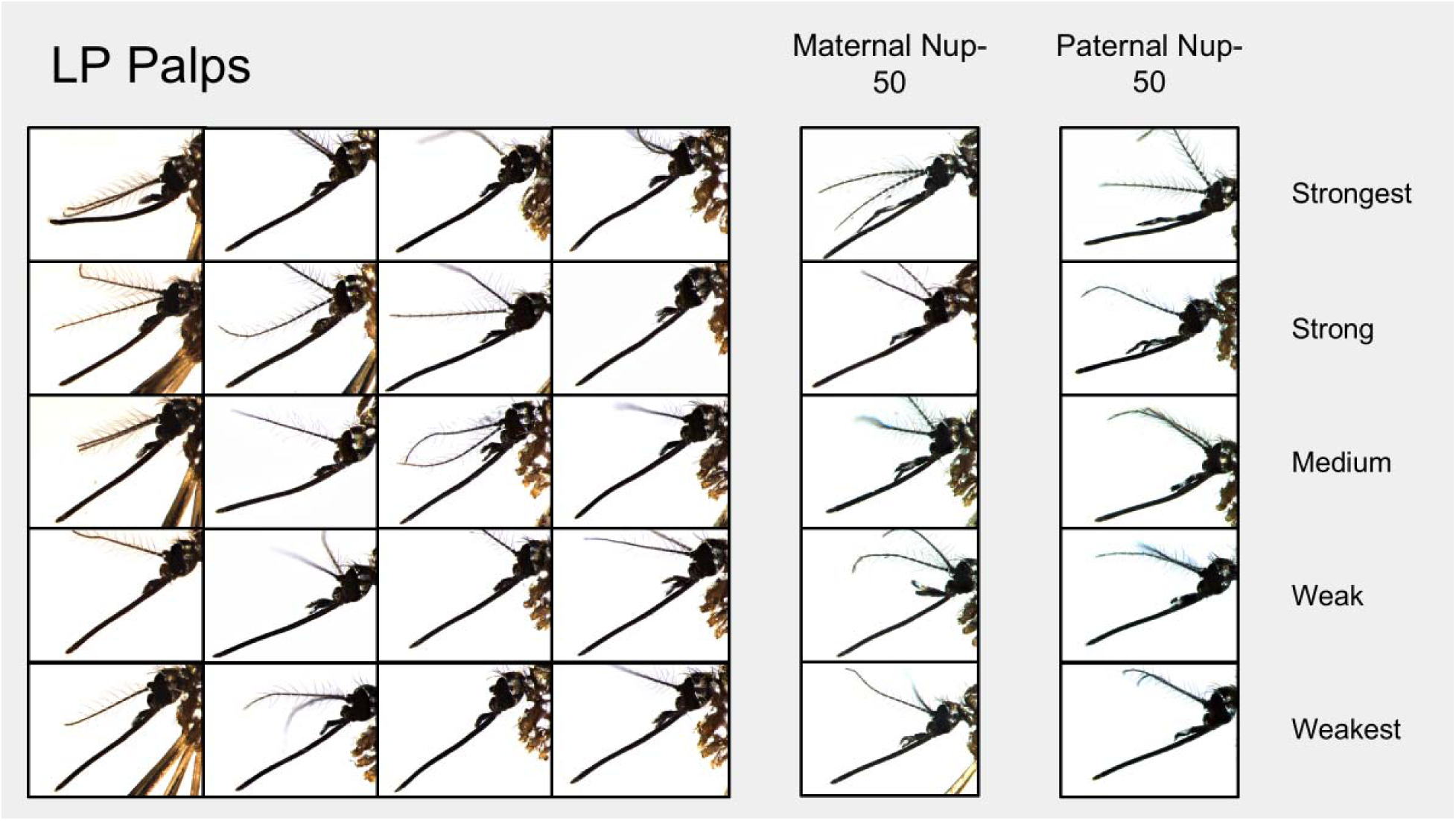
Palp abnormalities in female Nup50Cas9/GDe_dsx_ transheterozygotes. Representative palp morphologies of Liverpool wild-type females and female transheterozygotes carrying Nup50Cas9 and GDe_dsx_. Transheterozygous females generated with either maternally or paternally inherited Cas9 exhibited a range of palp abnormalities associated with androgenization. Representative phenotypes are arranged to illustrate increasing phenotypic severity.

**Supplementary Figure 7.**
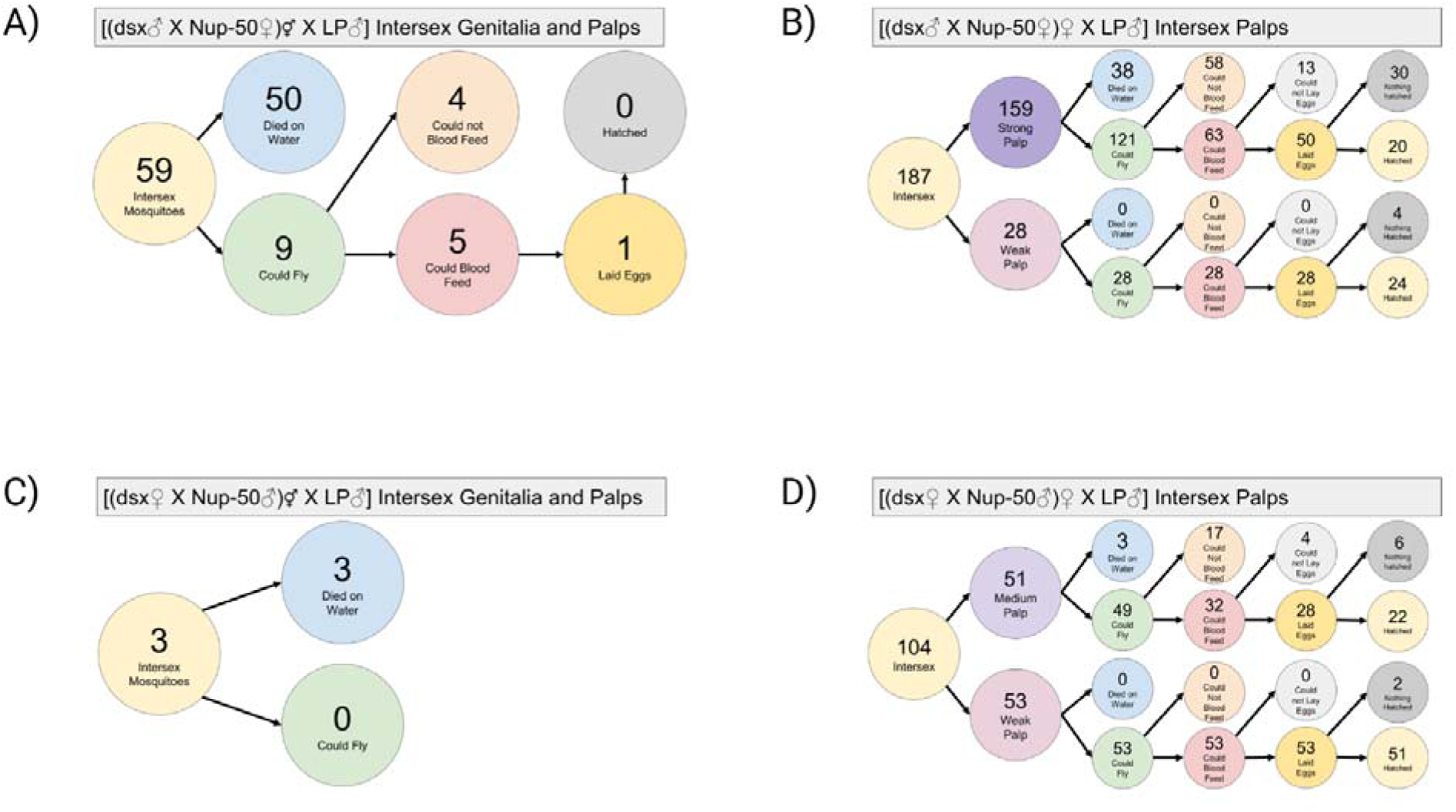
Survival and reproductive outcomes associated with androgenization in female offspring from GDedsx and Nup50Cas9 crosses. Female offspring from GDe_dsx_ and Nup50Cas9 crosses were phenotypically classified according to the severity of androgenization and followed through sequential developmental and reproductive milestones. Newly eclosed adults were first assessed for their ability to escape the water cup and fly. Surviving females were subsequently offered a blood meal, and blood-fed females were paired individually with wild-type males for assessment of fecundity and fertility. **(A–B)** Offspring generated from paternal GDe_dsx_ and maternal Nup50Cas9 crosses. **(A)** Individuals exhibiting strong androgenization of the genitalia during the pupal stage also displayed abnormal palp morphology. **(B)** Females classified as having weak or strong androgenization based primarily on palp morphology, as illustrated in **Supplementary** Figure 8, were followed through adult survival, blood feeding, oviposition, and egg hatch. **(C–D)** Offspring generated from maternal GDe_dsx_ and paternal Nup50Cas9 crosses. **(C)** Pupae exhibiting strong intersex genital phenotypes also displayed pronounced palp androgenization. **(D)** Females categorized according to palp phenotype were followed throughout adulthood to assess survival and reproductive performance.

**Supplementary Figure 8.**
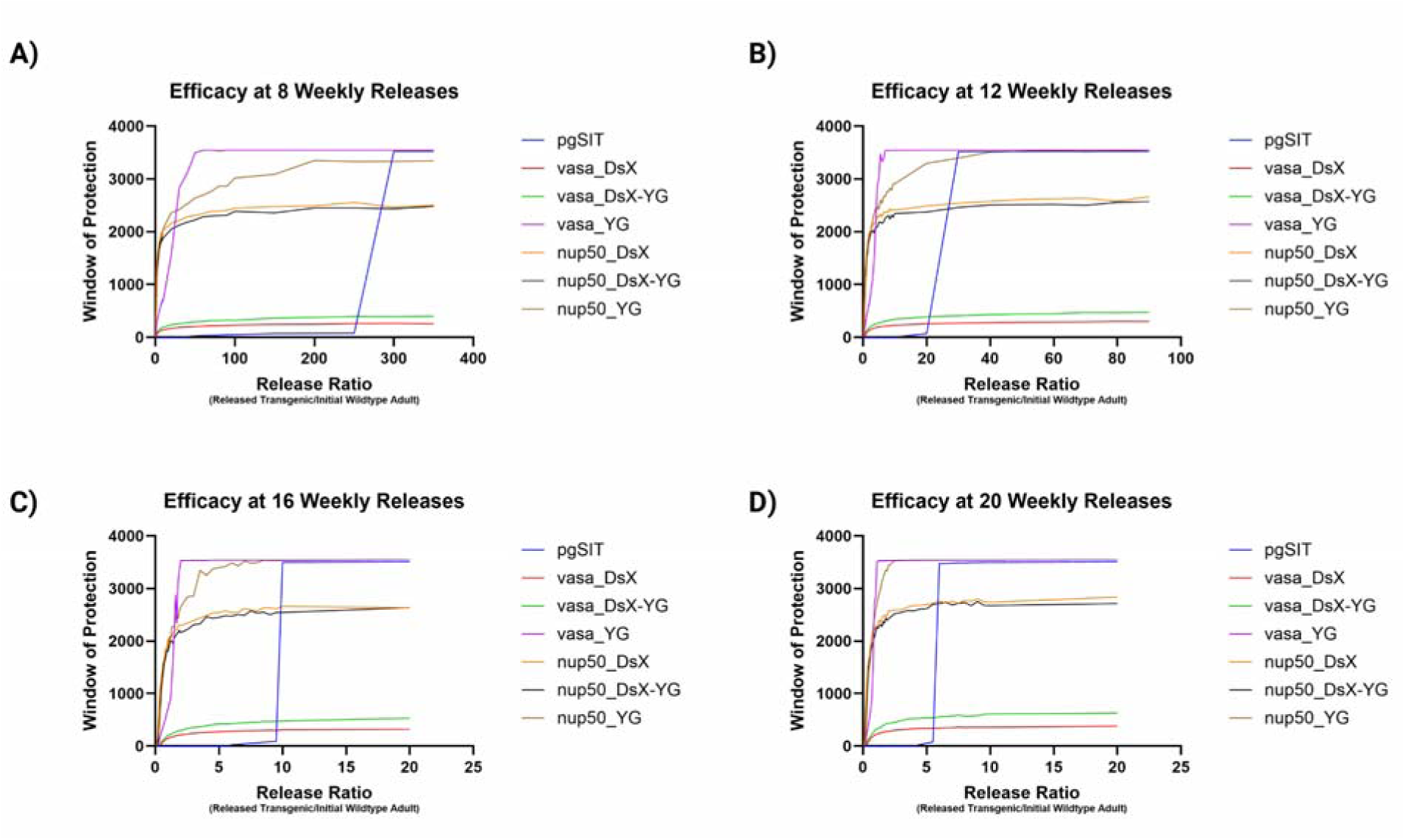
Predicted population suppression across release ratios and release durations. Experimentally parameterized split suppression drives, theoretical optimized-drive configurations, and pgSIT were compared across release programs lasting **(A)** 8, **(B)** 12, **(C)** 16, and **(D)** 20 weeks. For each release schedule, the predicted window of protection (WOP) is shown as a function of the adult male release ratio. WOP was defined as the duration for which the non-transgenic female population remained at least 90% below its pre-release equilibrium abundance in at least 50% of simulations. Simulations were run for 10 years; therefore, the maximum observable WOP was 3,650 days.

**Supplementary Figure 9.**
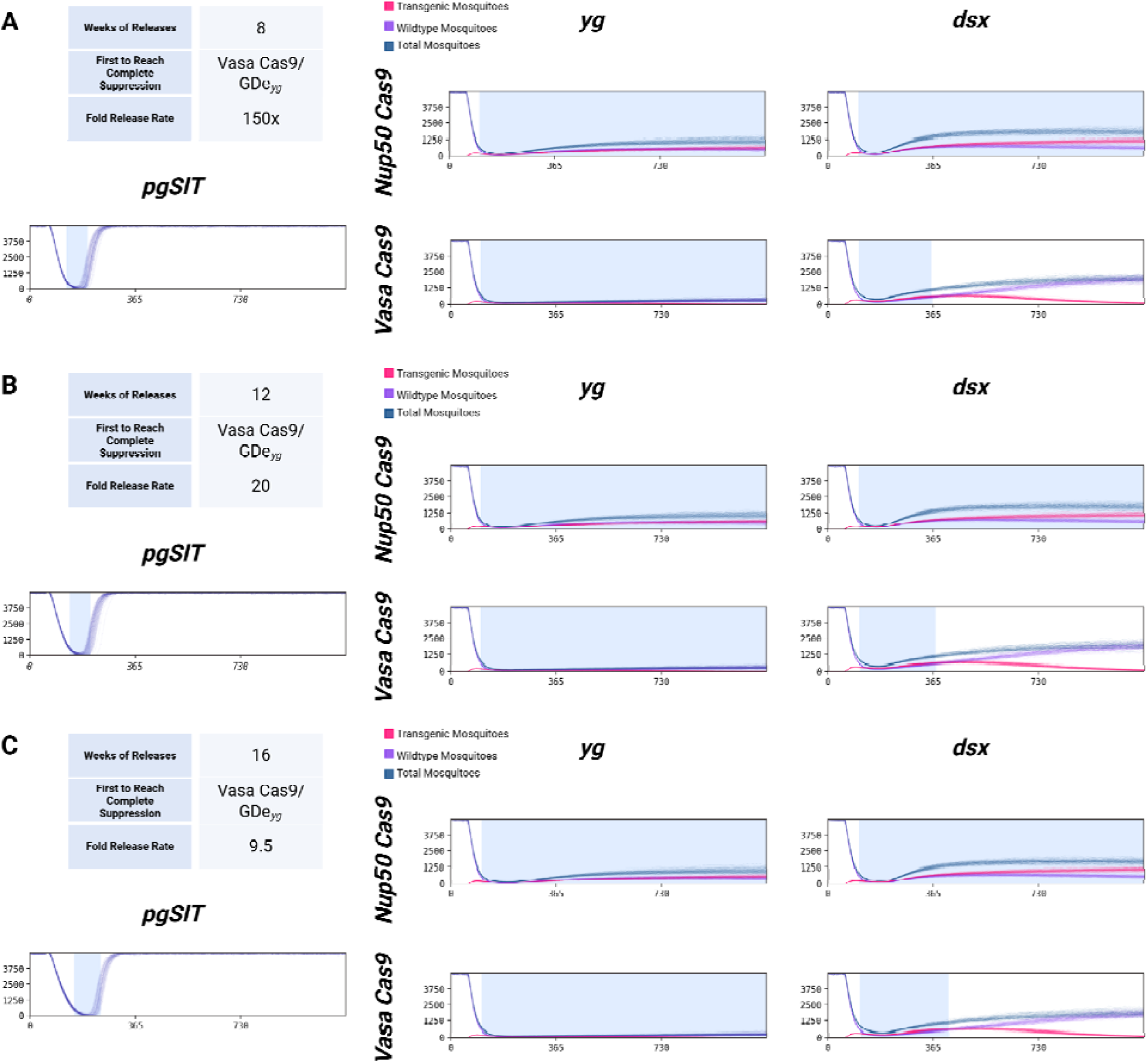
Population dynamics of individual suppression strategies at selected release schedules. Population trajectories for individual suppression strategies were compared using release conditions corresponding to the minimum pgSIT release ratio that produced the defined suppression criterion for each release duration. **(A)** For an 8-week release program, adult transgenic males were released at a 150:1 ratio relative to the initial wild population. **(B)** For a 12-week release program, males were released at a 20:1 ratio. **(C)** For a 16-week release program, males were released at a 9.5:1 ratio. The same release ratio was applied to each suppression strategy within a given panel to facilitate comparison. Simulations were initialized with 10,000 adult mosquitoes, releases began on day 50, and trajectories are displayed for 3 years. Blue shading denotes the window of protection (WOP), during which the non-transgenic female population met the predefined suppression threshold.

**Supplementary Figure 10.**
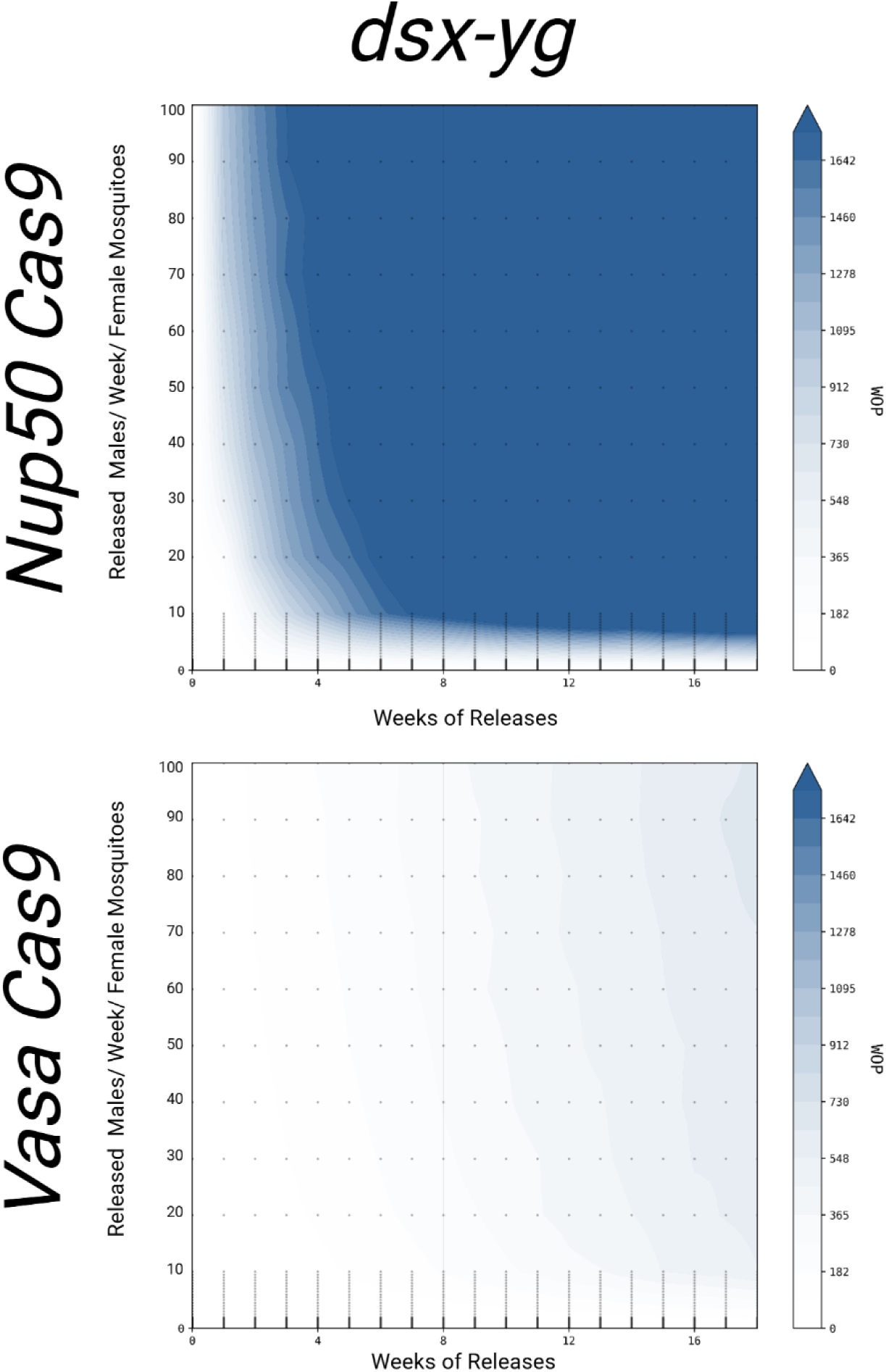
Modeling theoretical optimized split suppression drives combining GDe_yg_ inheritance and GDe_dsx_ reproductive effects. Theoretically optimized split-drive configurations were generated by combining the experimentally measured inheritance characteristics of the GDe_yg_ systems with the stronger suppression-associated reproductive fitness effects measured for the corresponding GDe_dsx_ systems. These hypothetical configurations were simulated using the same population-modeling framework and release conditions used for the experimentally parameterized split-drive systems in Figure 4. The analysis evaluates whether combining greater inheritance bias with stronger reproductive effects substantially improves predicted population suppression.

